# Novel function of Contactin associated protein 1 (Caspr 1)/ Paranodin in embryonic cortical neurons: hypoxia modulated neurite development

**DOI:** 10.1101/2025.03.02.641114

**Authors:** Gowthaman Suresh, Reshma Ramachandran, Sapana Sharma, Konstanze F Winklhofer, Vasudharani Devanathan

## Abstract

Hypoxia, a condition of inadequate oxygen supply, is a common phenomenon affecting neurons and brain tissue, leading to significant implications for neuronal health and function. The prevalence of hypoxia in the brain is associated with various neurological conditions, making it a critical area of study. Neuritogenesis, the process of neurite outgrowth, is an essential aspect of neuronal development and connectivity and is particularly sensitive to hypoxic stress. Investigating how hypoxia affects neurite outgrowth is vital for understanding neuronal response and adaptation under low oxygen conditions. This study explores how hypoxic stress affects neurite regulation mediated by Contactin Associated Protein-1 (Caspr1) in primary mouse embryonic cortical neurons. Hypoxia, induced by culturing neurons in a 2% oxygen environment, significantly reduced neurite length and induced notable changes in growth cone morphology. Concurrently, we observed an upregulation in the expression of Caspr1 and its transcriptional regulator C/EBPα, suggesting a compensatory role for Caspr1 in neurite extension under low oxygen conditions. Shorter hypoxia exposure periods revealed a dynamic biphasic response in Caspr1 levels, with an initial decrease followed by a substantial increase, correlating with corresponding changes in neurite length. This pattern emphasizes the critical involvement of Caspr1 in adapting neurite growth to fluctuating hypoxia duration. Furthermore, comparative analyses using wild-type and Caspr1 knockout Neuro2a cells demonstrated that the absence of Caspr1 mitigates hypoxia-induced neurite shortening, indicating a potential protective role against hypoxic stress. Additionally, hypoxia profoundly impacted mitochondrial morphology and function. Under hypoxic conditions, mitochondria transitioned to a more spherical shape. Mitochondrial respiration analysis revealed significant reductions in oxygen consumption rates (OCR), highlighting compromised mitochondrial function during hypoxia. These findings underscore the multifaceted role of Caspr1 in neurite regulation and mitochondrial adaptation to hypoxic stress. The study provides insights into the molecular mechanisms underpinning hypoxia-induced changes in neuronal morphology and function. Understanding these processes opens avenues for therapeutic strategies targeting Caspr1 in treating neurological disorders characterized by hypoxic stress. Future research will benefit from extending these investigations to more complex models, such as brain organoids, to further elucidate the metabolic and structural changes under hypoxia and their implications for neurodegenerative diseases.

## Introduction

Adequate oxygen supply is crucial for the normal physiological function of mammalian cells. Oxygen deprivation, or hypoxia, can lead to cellular dysfunction and, in severe cases, cell death (Kaelin & Ratcliffe, 2008). Metazoan cells have evolved sophisticated mechanisms to sense and respond to changes in oxygen availability, with the hypoxia-inducible factor pathway playing a central role in this process (Kaelin & Ratcliffe, 2008). Hypoxia can have profound effects on cellular metabolism, gene expression, and cell fate decisions (Al Tameemi et al., 2019; Nakayama & Kataoka, 2019).

Neurons, as highly specialized and energy-dependent cells, are particularly vulnerable to the detrimental effects of hypoxia. This vulnerability can lead to significant changes in the structure and function of the neurons. Brain hypoxia is not merely a rare pathological event; it can also occur in a healthy brain due to natural variations in oxygen levels across different brain regions. This condition may arise under normal physiological circumstances or during various diseases that disrupt the delivery and distribution of oxygen (Wallace et al., 2007). When hypoxia occurs, with oxygen levels dropping to 1%, it leads to the downregulation of genes critical for a wide range of neuronal activities. These include aspects such as neurotransmitter transportation, synaptic communication, and learning processes (H et al., 2011).

One key aspect of neuronal development is neurite outgrowth, which is essential for the formation of functional neural networks. It involves complex signaling pathways regulated by protein kinases, which control cytoskeletal rearrangements and membrane expansion (Bennison et al., 2020). The study of neuritogenesis is significant for understanding neuronal development and potential treatments for neurodegenerative diseases.

Hypoxia significantly affects neuronal morphology, with studies revealing various structural changes. In mouse striatal neurons, prolonged hypoxia led to decreased dendritic length without altering dendritic volume (Wallace et al., 2007). At the cellular level, hypoxia caused mitochondrial swelling, disorganization of cristae, and reduced cytoplasmic matrix density in neurons (Su, 1975). Presynaptic terminals and postsynaptic dendrites showed clearing and degeneration, while longer hypoxic periods resulted in extensive cytoplasmic rarefaction and organelle loss (Su, 1975). In rat somatosensory cortex neurons, oxygen deprivation significantly reduced axonal collaterals but did not affect dendritic proportions or spine density (Schröder & Luhmann, 1997).

Studying the impact of hypoxia on neurites is crucial due to its significant effects on neural development and function. Hypoxia can induce neurite outgrowth through adenosine A2A receptor activation, distinct from NGF-mediated pathways (O’Driscoll & Gorman, 2005). Chronic hypoxia leads to calcium overloading and oxidative stress, resulting in neurite retraction and neuronal damage (Saha et al., 2023). However, neuroprotective agents like erythropoietin can mitigate hypoxia-induced damage by promoting synaptogenesis and neurite repair, improving synaptic function and spatial memory (Xiong et al., 2019). Hypoxia disrupts CNS connectivity by altering axon pathfinding and synapse development, with HIF1α as the central mediator of the hypoxic response (Hofmeijer et al., 2014; Pocock & Hobert, 2008).

Given the profound influence of hypoxia on neurite dynamics, understanding the molecular players that regulate neurite outgrowth and connectivity becomes essential. Among these, the Contactin-associated protein-1 (Caspr1) is a cell adhesion molecule that has been implicated in the regulation of neurite outgrowth and neuronal connectivity. Caspr1 interacts with various signaling pathways and cytoskeletal components to modulate the dynamics of neurite formation and extension (Sultan et al., 2020). Under normal conditions, contactin-associated protein-1 inhibits neurite outgrowth and supports the establishment of a complex neuronal network (Devanathan et al., 2010). It plays a key role in maintaining axon-glial junctions, especially at paranodal regions of myelinated neurons (Einheber et al., 1997). Additionally, a family of Caspr1 proteins is increasingly linked to neurological disorders and neurodegenerative conditions. Dysregulation or mutations in Caspr family of proteins are associated with multiple sclerosis, marked by disrupted neuronal signaling due to demyelination (Coman et al., 2006; Wolswijk & Balesar, 2003), and autism spectrum disorders, involving altered synaptic connectivity (Anderson et al., n.d.; Falivelli et al., n.d.; Fernandes et al., n.d.). Caspr dysfunction may also contribute to axonal degeneration and impaired neuronal communication, making it a potential therapeutic target in diseases like Alzheimer’s and Parkinson’s.

Understanding the effects of hypoxia on neurite development and the role of Caspr1 in this process is crucial for unraveling the molecular mechanisms underlying neuronal adaptation and dysfunction in adverse conditions. Given that hypoxic stress is a common feature in various neurodevelopmental and neurodegenerative disorders, investigating how Caspr mediates neurite regulation under such conditions can provide valuable insights into the pathophysiology of these diseases. This knowledge not only enhances our understanding of neuronal resilience and vulnerability but also opens avenues for developing targeted therapeutic strategies to mitigate the impact of hypoxia on the developing and mature brain. In this paper, we have investigated Caspr-mediated neurite regulation in cortical neurons under hypoxic stress conditions.

## Materials and Methods

### Primary Neuron Culture

Embryonic cortical neurons were isolated from pregnant mice at day 16.5 using a modified protocol (Hilgenberg & Smith, 2007). Plates were pre-coated with 25 µg/mL poly-L-lysine and incubated overnight at 37°C. The following day, the coating was removed, and plates were washed with DPBS twice. Neuronal culture media (Neurobasal) was pre-warmed to 37°C. After the pregnant mouse was euthanized using CO2 and cervical dislocation, embryos were extracted and placed in ice-cold PBS. The embryos were decapitated, and cortical neurons were isolated. The brain was dissected out, meninges removed, and the cortex isolated and washed with HBSS. Cortices were enzymatically dissociated using acetylated trypsin (1 mg/100 µL) followed by DNaseI treatment (36 µL). Neurons were mechanically dissociated by triturating the tissue, and cells were filtered through a 70 µM cell strainer to remove clumps. Cell counts were performed using a hemocytometer, and neurons were plated at 20,000 cells/well for immunofluorescence in a 24-well plate or 200,000 cells/well for protein and RNA isolation in a 6-well plate. Growth media was replaced after 2 hours, and half of the media was changed every three days.

### Cell Viability Assay

Cortical neurons were cultured for 3-4 days in a 24-well plate, followed by hypoxia or normoxia treatments for 24 hours. Cell viability was assessed using the CCK-8/WST-8 assay, which detects the reduction of tetrazolium salt by cellular dehydrogenases to orange formazan. The absorbance was measured at 460 nm using the Cytation5 instrument.

### Neuro2a Cell Culture

Neuro2a cells were obtained from ATCC and cultured in DMEM supplemented with 10% fetal bovine serum and 1% antibiotics. Cells were maintained at 37°C in 5% CO2 and 90% humidity. Once 70% confluence was reached, cells were trypsinized, counted, and reseeded for the experiments.

### Protein Isolation and Western Blotting

Total protein was isolated either through direct cell lysis or using lysis buffer. For direct lysis, cells were scraped from culture dishes after PBS washes and lysed in 2x Laemmli sample buffer (LSB). Protein was denatured at 95°C for 10 minutes. Alternatively, cells were lysed using a buffer followed by centrifugation, and protein quantification was performed using a BCA assay. Proteins were separated using SDS-PAGE and transferred to nitrocellulose membranes. Membranes were blocked with 5% bovine serum albumin or skimmed milk and incubated with primary antibodies overnight at 4°C. HRP-linked secondary antibodies were added, and chemiluminescence was detected using a Biozyme system or Azure imager.

### Immunostaining

Cells cultured on coverslips were fixed with 4% paraformaldehyde and permeabilized with 0.2% Triton X-100. After blocking with 5% normal goat serum, cells were incubated with primary antibodies overnight at 4°C, followed by secondary antibodies at room temperature. Nuclei were stained with DAPI, and coverslips were mounted for microscopy.

### Microscopy

Structured illumination microscopy (SIM) or laser scanning microscopy (LSM) was conducted using a Zeiss ELYRA PS.1 LSM880 microscope. Images were acquired at 20x or 63x magnification using a 1.4 oil-immersion objective. SIM was performed with five phases and three to five rotations of the SIM grid. Imaging settings were standardized within experiments, and raw images were processed using Zeiss ZEN software.

### 3D Reconstruction of Mitochondria

Mitochondria were imaged using SIM and reconstructed in 3D using Imaris 10.0 software. Z-stacks were taken at intervals of 0.144 µm, and mitochondrial surfaces were reconstructed. Quantitative data were extracted and analyzed using GraphPad Prism.

### Seahorse Mito Stress Assay

Mitochondrial respiration was evaluated using the Seahorse XFe96 Analyzer. Cortical neurons were seeded at 20,000 cells per well and cultured for 4-5 days before treatment with 200 µM cobalt chloride to induce hypoxia. Cells were washed with Seahorse XF media and treated with oligomycin (1 µM), FCCP (0.5 µM), and a combination of rotenone and antimycin A (0.5 µM each). Oxygen consumption rates (OCR) were measured in real-time, and data were analyzed using Agilent Wave Pro software.

### RNA Isolation and RT-PCR

Total RNA was isolated from the cells using a TRIzol reagent. 1 μg of total RNA was used for cDNA synthesis by iScript cDNA Synthesis Kit (Bio-Rad) or SuperScript III First-strand cDNA Synthesis Supermix (Invitrogen). Quantitative PCR was performed using 100 ng of cDNA, iTaq Universal SYBR Green Supermix (Bio-Rad) and gene-specific primers in Bio-Rad CFX96/384 Real-Time PCR detection system for 40 cycles with denaturation (10 sec, 95°C) and annealing/extension (30 sec, 58°C). β-actin was used as a housekeeping gene to normalize the mRNA expression levels of all the genes.

### Neurite Length Measurement

Neurite outgrowth was quantified using ImageJ software and the NeuronJ plugin. Fixed neurons were stained with β-III tubulin and imaged at 10x or 20x magnification. Neurite lengths were measured from random fields of view, and the data were analyzed accordingly.

### Cloning of sgRNAs into pU6-(BbsI)_CBh-Cas9-T2A-mCherry for Caspr1 Knockout

To generate a Caspr1 knockout, two sgRNAs targeting exons 4 and 5 of the Caspr1 gene were designed using crispr.mit.edu, with a preference for those exhibiting low mismatch scores and high target specificity. The pU6-(BbsI)_CBh-Cas9-T2A-mCherry plasmid was digested using BbsI enzyme, and the sgRNAs were phosphorylated with T4 polynucleotide kinase before cloning into the vector. The sgRNA-containing plasmids were then transformed into DH5α *E. coli* cells and selected on ampicillin-containing agar plates. Positive clones were identified by colony PCR, confirmed by Sanger sequencing, and the plasmids were extracted using the Nucleospin Plasmid Miniprep kit.

The sgRNAs were assembled into a single plasmid using Gibson assembly. The reaction was carried out at 50°C for 1 hour, followed by transformation into DH5α *E. coli*. Positive clones were verified through colony PCR using Gibson-specific primers and confirmed by sequencing. Plasmid isolation was performed with the Qiagen Midi Prep kit.

Neuro2a cells were transfected with the Caspr1-targeting plasmid using Lipofectamine 3000, following the manufacturer’s instructions. Post-transfection, mCherry expression confirmed successful transfection. Cells were cultured in 96-well plates at a density of one cell per well for two weeks and screened via PCR and Western blot. Primers flanking the target region were used for PCR, while an anti-Caspr1 antibody was employed for Western blot detection.

### Genomic DNA Isolation from Neuro2a Cells

Genomic DNA was extracted from transfected Neuro2a cells cultured to 90% confluency. Cells were harvested by trypsinization and lysed using Proteinase K and buffer AL at 56°C for 10 minutes. Ethanol was added, and the lysate was passed through a DNeasy mini spin column (Qiagen) to bind genomic DNA. The column was washed with buffers AW1 and AW2, followed by elution with nuclease-free water.

### Cloning of Mouse Caspr1 Promoter into pGL2 Basic Vector

The mouse Caspr1 promoter was cloned into the pGL2 basic vector. Primers containing SmaI and XhoI restriction sites were designed to amplify the Caspr1 promoter from Neuro2a genomic DNA. PCR amplification was carried out for 35 cycles using Pfu polymerase. The PCR product was separated via agarose gel electrophoresis, and the desired band was extracted using the Takara gel extraction kit. The pGL2 vector and promoter insert were digested with SmaI and XhoI and ligated overnight at 16°C using T4 DNA ligase. Following transformation into DH5α *E. coli* and plating on ampicillin-LB-agar, positive clones were identified by restriction digestion and Sanger sequencing.

### Plasmid Isolation from *E. coli* Using Miniprep and Midiprep Kits

Plasmids were isolated from *E. coli* using the Nucleospin Plasmid Miniprep kit (Takara) for small-scale preparations and the Qiagen Midiprep kit for larger-scale extractions. For minipreps, 5 ml of overnight *E. coli* cultures were harvested and processed following the kit protocol, involving cell lysis, neutralization, and purification using a spin column. For midipreps, 50 ml cultures were processed by binding plasmid DNA to a resin column, washing, and eluting in nuclease-free water after precipitation with isopropanol.

### Agarose Gel Electrophoresis

Agarose gel electrophoresis was used to separate DNA fragments. A 1% agarose gel was prepared in 1X TAE buffer containing ethidium bromide. DNA samples were loaded and electrophoresed in 1X TAE running buffer. After electrophoresis, DNA bands were visualized under UV light.

### Plasmid Transfection into Neuro2a Cells

Neuro2a cells were transfected with plasmid DNA using Lipofectamine 3000. Cells were seeded in 6-well plates in antibiotic-free DMEM medium and transfected at 40-50% confluency with a Lipofectamine-DNA mixture prepared in Opti-MEM. After 6 hours of incubation, the medium was replaced with complete DMEM. Post-transfection, cells were harvested for RNA, DNA, or protein extraction after 24 hours.

### Reporter Assays

Luciferase reporter assays were performed 24 hours post-transfection using the Promega Luciferase Assay System. Neuro2a cells were lysed in 1X reporter lysis buffer, and luciferase activity was measured in a luminometer. Luciferase activity was normalized to β-galactosidase activity, determined using the β-galactosidase Enzyme Assay System (Promega) at 420 nm.

## Statistical Analysis

GraphPad Prism 9.5 and Microsoft Excel were used for statistical analyses. Data are presented as mean ± SD. Statistical significance was evaluated using the Student’s t-test (for comparisons between two groups), or ANOVA (for comparisons between more than two groups), with significance levels, indicated as *P < 0.05, **P < 0.01, and ***P < 0.001.

## Results

### Effects of Hypoxia on Neuronal Viability and Morphology

Hypoxia, defined by the absence of adequate oxygen in cells, poses a significant threat to neurons due to their high oxygen dependency. To assess the impact of hypoxia on neuronal morphology, we first evaluated cell viability under hypoxic conditions using the CCK-8 assay. The results showed that hypoxia did not compromise neuronal viability, as cell survival rates remained consistent between hypoxic and normoxic conditions.

Next, we examined the morphological alterations in neurons after 24 hours of hypoxic exposure. Bright-field imaging revealed that neurons under hypoxia exhibited notable morphological changes, including a significant reduction in neurite length compared to neurons maintained under normoxia. To confirm the induction of hypoxia, we performed a Western blot analysis for Hypoxia-inducible factor 1-alpha (HIF1α), a well-established marker of cellular adaptation to hypoxic stress. It is stabilized and activated under low oxygen conditions. However, under normoxic conditions, HIF1α is rapidly degraded through the ubiquitin-proteasome pathway, mediated by oxygen-dependent prolyl hydroxylation. This degradation prevents its accumulation and transcriptional activity, ensuring that hypoxia-responsive genes remain inactive in the presence of sufficient oxygen. However, during hypoxic conditions, the activity of prolyl hydroxylases is suppressed due to the lack of oxygen, preventing hydroxylation. The tightly regulated expression of HIF1α highlights its role as a dynamic sensor of oxygen levels. The results confirmed the hypoxic state in treated neurons by showing elevated HIF1α protein levels.

To quantitatively assess the impact of hypoxia on neurite structure, neurons were fixed, stained, and analyzed using the NeuronJ plugin in FIJI ImageJ. The analysis demonstrated a significant decrease in neurite length in hypoxia-treated neurons compared to those in normoxic conditions.

Given the detrimental effect of hypoxia on neurite length, we investigated the potential for recovery upon return to normoxic conditions. Neurons were subjected to 24 hours of hypoxia, followed by a return to normoxia for an additional 24 hours. Time-lapse imaging at 0, 3-, 6-, 12-, and 24-hours post-hypoxia revealed that the neurite length, which had decreased due to hypoxic stress, was restored after 24 hours in normoxia. This suggests that the structural damage induced by hypoxia is reversible upon reoxygenation.

**Figure 1:**
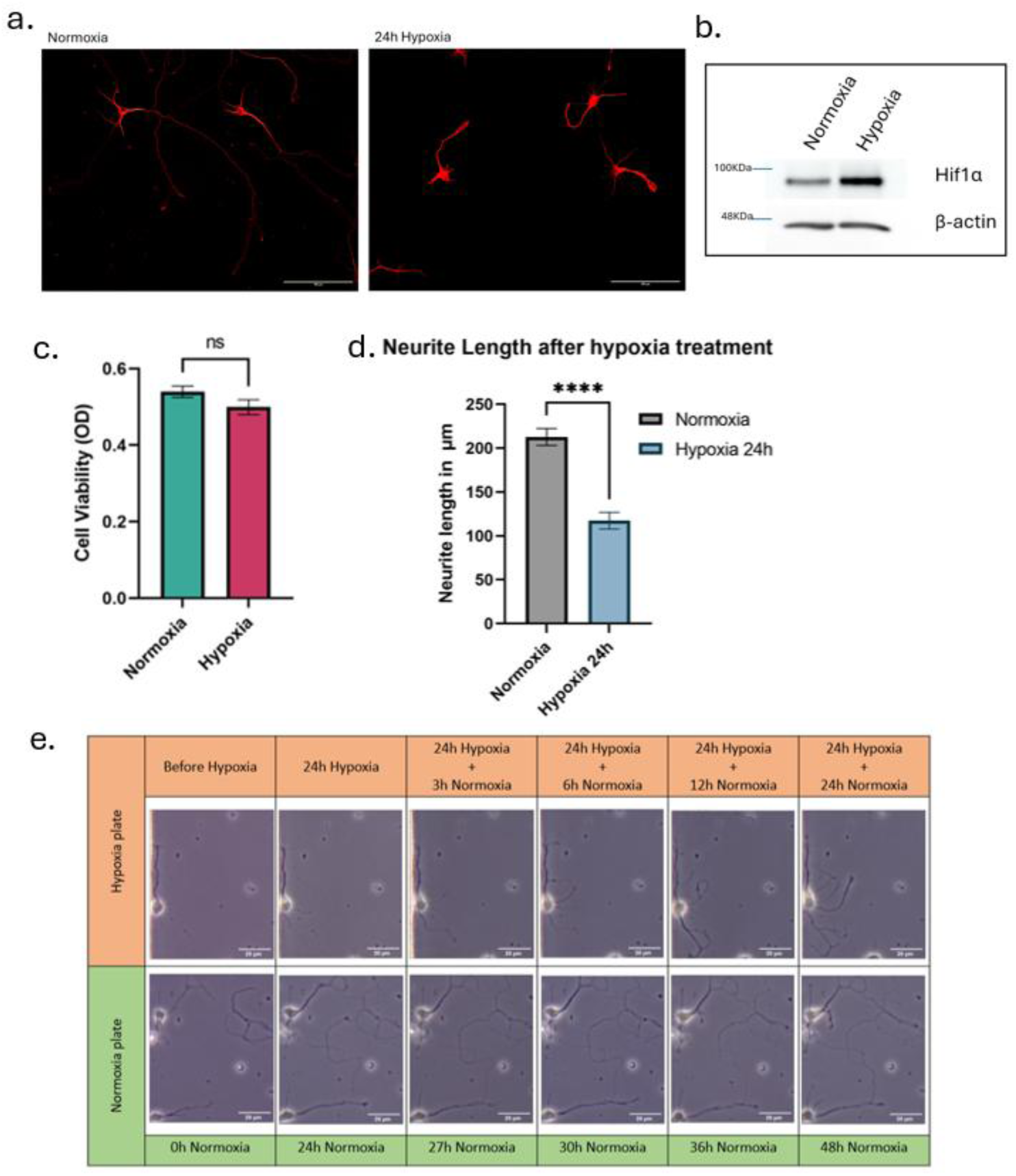
Hypoxia-Induced Alterations in Neuronal Morphology, Viability, and Protein Expression: (a) Bright-field images depict significant morphological changes in neurons after 24 hours of hypoxia compared to normoxic conditions. (b) Western blot analysis reveals elevated HIF1α expression in neurons subjected to hypoxia, indicating a cellular response to oxygen deprivation. (c) Bar graph comparing neuronal viability under normoxic and hypoxic conditions over a 24-hour period, showing no significant reduction in cell viability despite oxygen deprivation. N=3. Statistical analysis was performed using an unpaired t-test. p-value > 0.05 not significant (ns). (d) A bar graph with mean ± SEM illustrating the statistically significant reduction in neurite length following hypoxic treatment, underscoring the detrimental effects of oxygen deprivation on the structure of cortical neurons (N=5). Statistical analysis was performed using an unpaired t-test ****P<0.0001, evaluated using a two-tailed test. (e) The tile image (20X) sequence that demonstrates the impact of 24-hour hypoxia on neurite length and the subsequent recovery observed when neurons are returned to normoxic conditions for an additional 24 hours. The images collectively highlight the potential for reversibility of hypoxia-induced structural damage in neurons. This figure demonstrates the growth of neurite without any interruption in normoxia state (Control) for the overall period of experimental time.

### Hypoxia-Induced Regulation of Caspr1 and Neurite-Related Proteins

Previous studies have identified Caspr1 as a key cell adhesion molecule involved in the regulation of neurite length. Given our findings of reduced neurite length under hypoxic conditions, we investigated whether hypoxia influences Caspr1 expression. Neurons subjected to 24 hours of hypoxia were lysed to extract total protein, and subsequent Western blot analysis revealed a significant increase in Caspr1 protein levels in hypoxia-treated cells.

In parallel, quantitative PCR analysis of RNA extracted from these neurons demonstrated a significant increase in Caspr1 mRNA levels under hypoxic conditions, corresponding with the protein expression changes observed. To explore the transcriptional regulation of Caspr1 under hypoxia, we conducted a promoter activity assay. The Caspr1 promoter was cloned upstream of a luciferase reporter in the pGL2 plasmid and transfected into Neuro2a cells, which were then subjected to hypoxia. The assay revealed that Caspr1 promoter activity was significantly elevated under hypoxic conditions.

Bioinformatics analysis using Alibaba 2.0 identified three transcription factors—NF-1, Sp1, and C/EBPα—as potential regulators of the Caspr1 promoter. To confirm this, we overexpressed CEBPα in Neuro2a cells, which resulted in increased Caspr1 mRNA levels, indicating that CEBPα directly regulates Caspr1 transcription. This was further supported by Western blot analysis, which showed elevated CEBPα protein levels in cells under hypoxia.

In addition to Caspr1, we examined the expression levels of other key proteins involved in neurite regulation, including the Prion protein and RhoA. Western blot analysis revealed that hypoxia also altered the expression patterns of these proteins, suggesting that hypoxia exerts a broad influence on the molecular pathways that regulate neurite outgrowth.

**Figure 2:**
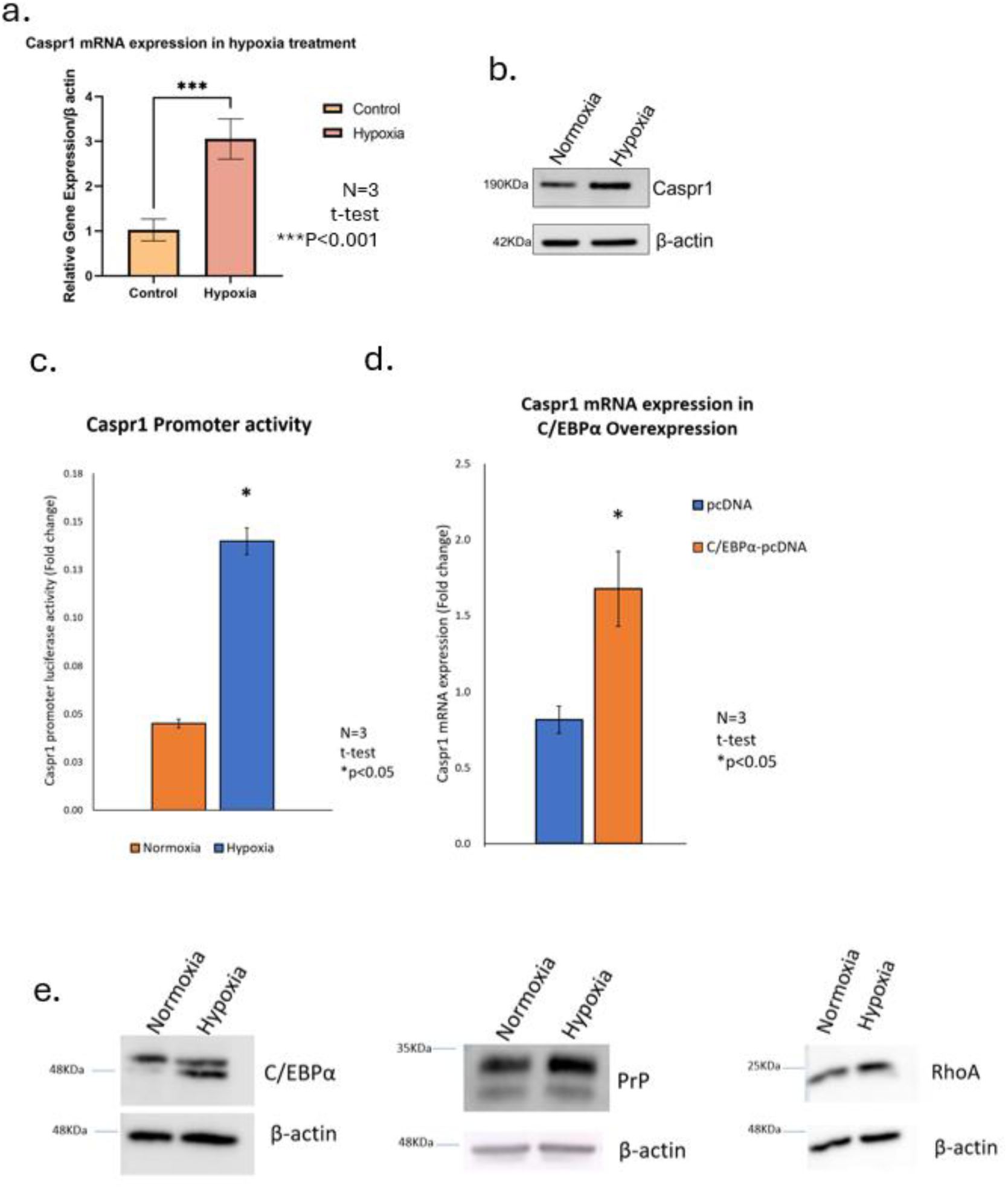
Hypoxia-Induced Regulation of Caspr1 Expression and Associated Proteins in Neurons. (a) A graphical quantification shows the mean ± SEM of quantitative mRNA expression levels of Caspr1 as determined by qRT-PCR, providing insights into the transcriptional regulation of Caspr1 under hypoxic stress (N=3). Statistical analysis was performed using an unpaired t-test. ***P<0.001, evaluated using a two-tailed test. (b) Western blot analysis showing differential Caspr1 protein levels in neurons exposed to hypoxia compared to normoxia, indicating changes at the protein level. (c) A bar graph shows the mean ± SEM of the activity of the Caspr1 promoter, measured using a luciferase assay under normoxic and hypoxic conditions. Statistical analysis was performed using an unpaired t-test. *P<0.05. (d) A bar graph with mean ± SEM depicting the changes in Caspr1 mRNA expression in control cells (pcDNA) versus cells overexpressing C/EBPα (C/EBPα-pcDNA). Statistical analysis was performed using an unpaired t-test. *P<0.05. (e) Western blot analysis of C/EBPα protein levels in neurons under normoxic and hypoxic conditions, underscoring its regulatory role in Caspr1 expression. Additionally, the figure presents Western blot analyses of neurite-regulating proteins, including Prion protein and RhoA, under hypoxic and normoxic conditions, illustrating how hypoxia differentially affects proteins critical for neurite outgrowth and stability.

### Impact of Hypoxia on Growth Cone Morphology and Caspr1 Localization

Given the significant effect of hypoxia on neurite length, we further examined the morphology of the growth cone—the neurite’s guidance structure—under hypoxic conditions. Neurons subjected to 24 hours of hypoxia were fixed, immunostained with the growth cone marker GAP43, and analyzed via microscopy. The results revealed that hypoxia induced a marked reduction in growth cone size, with growth cones appearing shrunken compared to those in normoxia-grown neurons. Quantitative analysis of growth cone surface area using ImageJ confirmed a significant decrease in size in hypoxia-treated neurons.

Considering the observed increase in Caspr1 expression under hypoxia and the concomitant collapse of growth cones, we investigated the localization of Caspr1 within the growth cones following hypoxic stress. Immunostaining for Caspr1 followed by microscopic analysis revealed a pronounced increase in Caspr1 expression within the growth cones of hypoxia-treated neurons, particularly at the growth cone tips, compared to neurons maintained under normoxic conditions. Quantitative analysis of mean fluorescence intensity further confirmed a significant elevation in Caspr1 expression at the tips of growth cones in hypoxia-treated neurons, implicating Caspr1 in the hypoxia-induced morphological alterations observed in growth cones. This implies a potential role of Caspr1 in the growth cone dynamics and morphological alterations observed during hypoxic stress, possibly contributing to the reorganization of cytoskeletal elements or influencing the signaling pathways that regulate neurite outgrowth. Additionally, the increased expression at the tips could suggest that Caspr1 plays a crucial role in the stabilization or destabilization of growth cone structures under adverse conditions.

**Figure 3:**
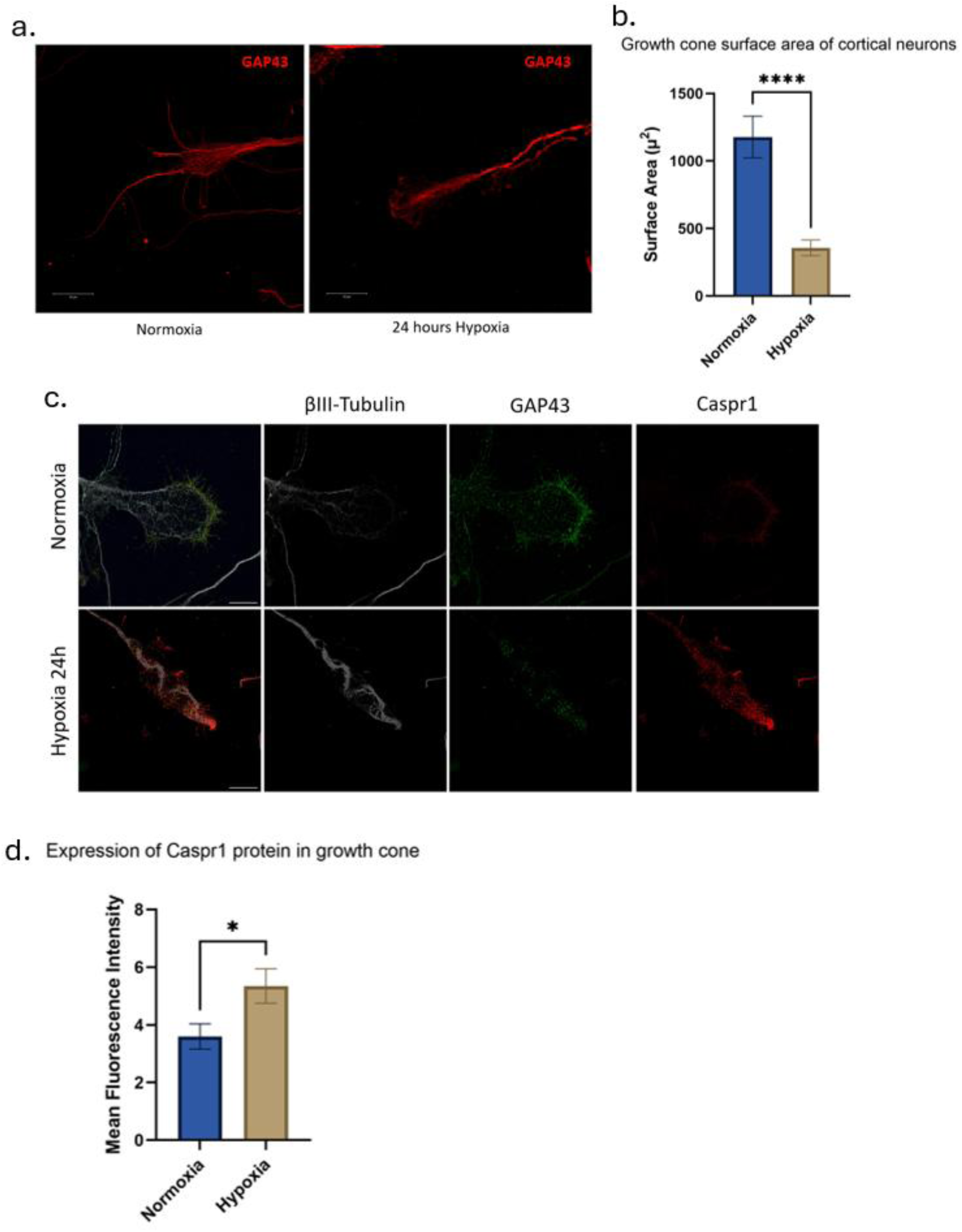
Hypoxia-Induced Alterations in Growth Cone Morphology and Caspr1 Expression. (a) Immunofluorescence images of growth cones stained with GAP43, comparing the morphology under normoxic and hypoxic conditions, and highlighting the structural changes induced by hypoxia. (b) A graph shows mean ± SEM quantifying the surface area of growth cones under both conditions, providing a measure of the extent to which hypoxia disrupts growth cone architecture (N=3). Statistical analysis was performed using an unpaired t-test. ****P<0.0001, evaluated using a two-tailed test. (c) Immunostained images of growth cones under normoxic and hypoxic conditions, with separate panels for βIII-tubulin, GAP43, and Caspr1, showcasing the differential expression and distribution of these proteins. (d) A bar shows mean ± SEM quantifying Caspr1 expression in the growth cone tips, illustrating the modulation of Caspr1 levels in response to hypoxic stress (N=3). Statistical analysis was performed using an unpaired t-test. *P<0.05, evaluated using a two-tailed test.

### Temporal Effects of Hypoxia on Neurite Length and Caspr1 Expression

Building on our previous findings that 24-hour hypoxia significantly reduces neurite length in cultured neurons, we explored the effects of shorter hypoxic durations. Cortical neurons were subjected to hypoxia for 3, 6, 12, and 24 hours, after which they were fixed, stained with the neurite marker βIII-Tubulin, and imaged via microscopy. Neurite length was quantified using the NeuronJ plugin in ImageJ software.

The analysis revealed a biphasic response in neurite length: an initial increase during shorter hypoxic periods (3 and 6 hours), followed by a significant reduction during prolonged hypoxia (12 and 24 hours). Additionally, neurons exposed to prolonged hypoxia (12 and 24 hours) exhibited not only reduced neurite length but also notable breaks along the neurites, indicating structural damage. The initial increase in neurite length during shorter hypoxic periods could be attributed to an adaptive response where neurons attempt to extend their neurites to seek out a more favorable environment. This adaptive mechanism might involve transient changes in the cytoskeletal architecture, enhancing neurite outgrowth initially. However, as the hypoxic condition prolongs, the cellular stress intensifies. Prolonged hypoxia leads to decreased ATP production, impairing cellular energy homeostasis. This energy deficit disrupts the functioning of the molecular motors and scaffolding proteins essential for maintaining and extending neurites. Additionally, the sustained hypoxic stress triggers cellular signaling pathways, including those mediated by hypoxia-inducible factors (HIFs), that lead to cytoskeletal reorganization, breakdown of actin filaments, and subsequent neurite retraction. Hence, this significant reduction in neurite length under prolonged hypoxia highlights the physiological limitations of neurons to cope with extended hypoxic conditions.

Given these temporal changes in neurite morphology, we investigated the expression of Caspr1 protein under varying durations of hypoxic stress. Western blot analysis of proteins extracted from hypoxia-treated neurons showed that Caspr1 expression was initially reduced at 3 and 6 hours of hypoxia but significantly increased at 12 and 24 hours. This pattern of Caspr1 expression closely mirrored the changes in neurite length, suggesting that Caspr1 is dynamically regulated in response to hypoxic stress, with its expression decreasing during early hypoxic exposure and increasing as hypoxia is prolonged.

**Figure 4:**
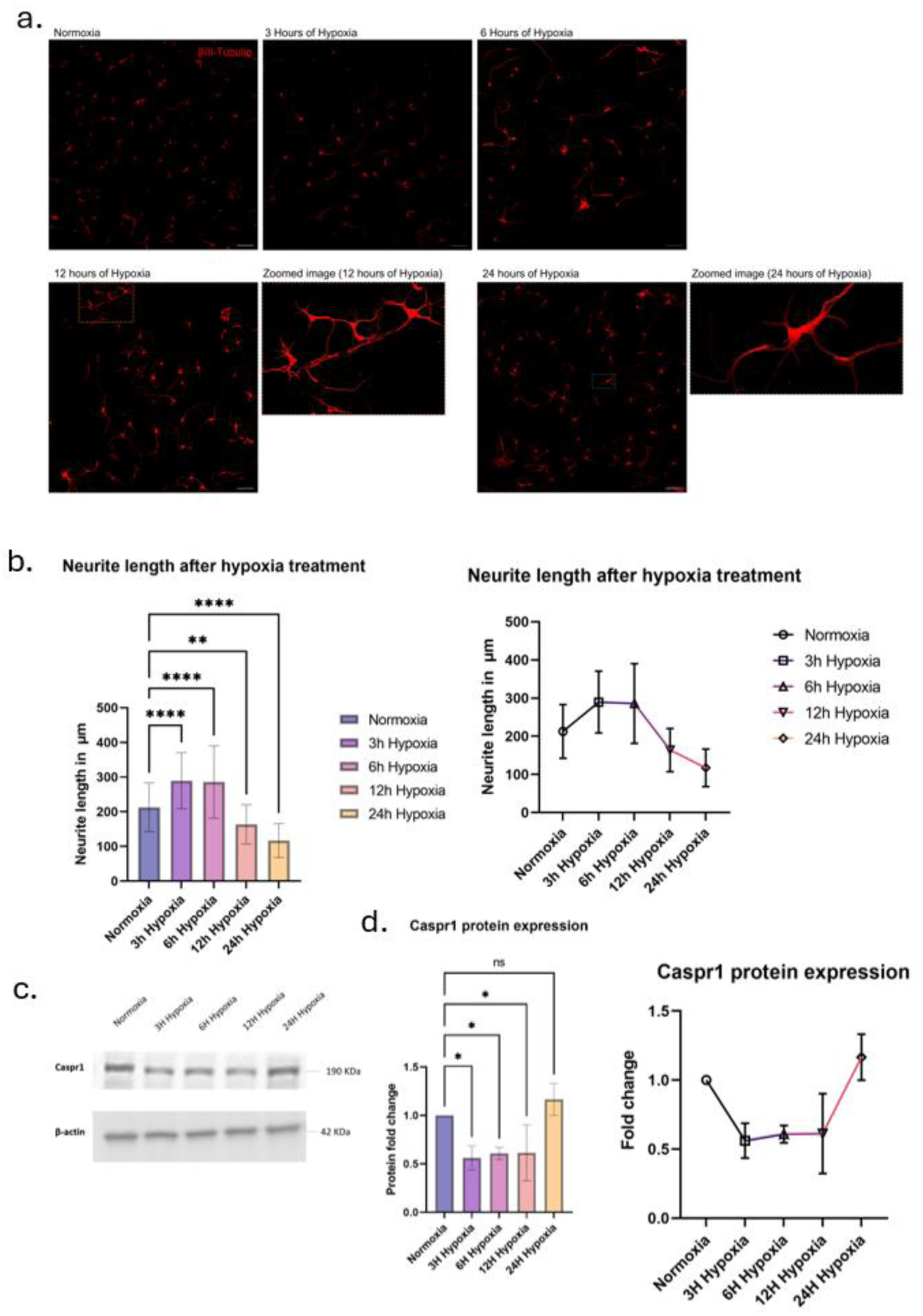
Temporal Dynamics of Neurite Outgrowth and Caspr1 Expression Under Hypoxic Conditions. (a) Representative immunofluorescence images of neurons stained with βIII-Tubulin, illustrating neurite length at various time points during hypoxia. (b) The graph shows mean ± SEM summarizing neurite length changes over short to prolonged periods of hypoxia, indicating the temporal dynamics of neurite outgrowth in response to oxygen deprivation (N=3). Statistical analysis was performed using ordinary One-way ANOVA with Dunnett’s multiple comparisons test, comparing the effects of time on neurite length. The degree of significance is represented as follows: ****P<0.0001 and **P<0.01. (c) Western blot analysis showing the expression pattern of Caspr1 protein across different durations of hypoxic stress, demonstrating how hypoxia affects Caspr1 over time. (d) A bar graph depicting mean ± SD of the fold change in Caspr1 protein levels over time under hypoxic conditions (N=3), offering a temporal view of Caspr1 regulation in response to hypoxia. Statistical analysis was performed using ordinary One-way ANOVA with Dunnett’s multiple comparisons test. The degree of significance is represented as follows: *P<0.05.

### Role of Caspr1 in Neurite Regulation and Response to Hypoxia

To further elucidate Caspr1’s role in neurite regulation, we attempted to generate a Caspr1 knockout (KO) Neuro2a cell line. Although efforts to create a homozygous Caspr1 KO line were not successful, we established a heterozygous Caspr1 knockout (ΔCaspr1) cell line, which exhibited a pronounced phenotype characterized by reduced neurite length. Therefore, the ΔCaspr1 line was utilized for subsequent experiments.

Live cell imaging comparing neurite outgrowth between wild-type and ΔCaspr1 Neuro2a cells revealed that ΔCaspr1 cells developed significantly longer neurites than their wild-type counterparts, underscoring Caspr1’s role as a negative regulator of neuritogenesis.

To further investigate, we assessed the expression of other key neurite-regulating factors— Prion protein, GAP43, and Rac1—in both wild-type and ΔCaspr1 Neuro2a cells under hypoxic conditions. After 24 hours of hypoxia, mRNA was extracted, converted to cDNA, and analyzed using qRT-PCR. The expression profiles of these factors were similar between wild-type and ΔCaspr1 cells, suggesting that Caspr1 exerts a more significant inhibitory effect on neurite length than these other factors.

Building on prior findings that Caspr1 loss leads to increased neurite length under normoxic conditions, we examined how ΔCaspr1 cells respond to hypoxic stress. Both wild-type and ΔCaspr1 Neuro2a cells were subjected to 24 hours of hypoxia, followed by fixation and neurite length measurement using the NeuronJ plugin in FIJI software.

Remarkably, ΔCaspr1 cells maintained significantly longer neurites under hypoxic conditions compared to wild-type cells. Moreover, the neurite lengths of ΔCaspr1 cells in hypoxia were comparable to those in normoxia, and were notably longer than the neurites of wild-type cells under normoxia. These findings suggest that the absence of Caspr1 may confer a protective effect against the morphological impacts of hypoxia, potentially offering resilience to hypoxia-induced neurite shortening.

**Figure 5:**
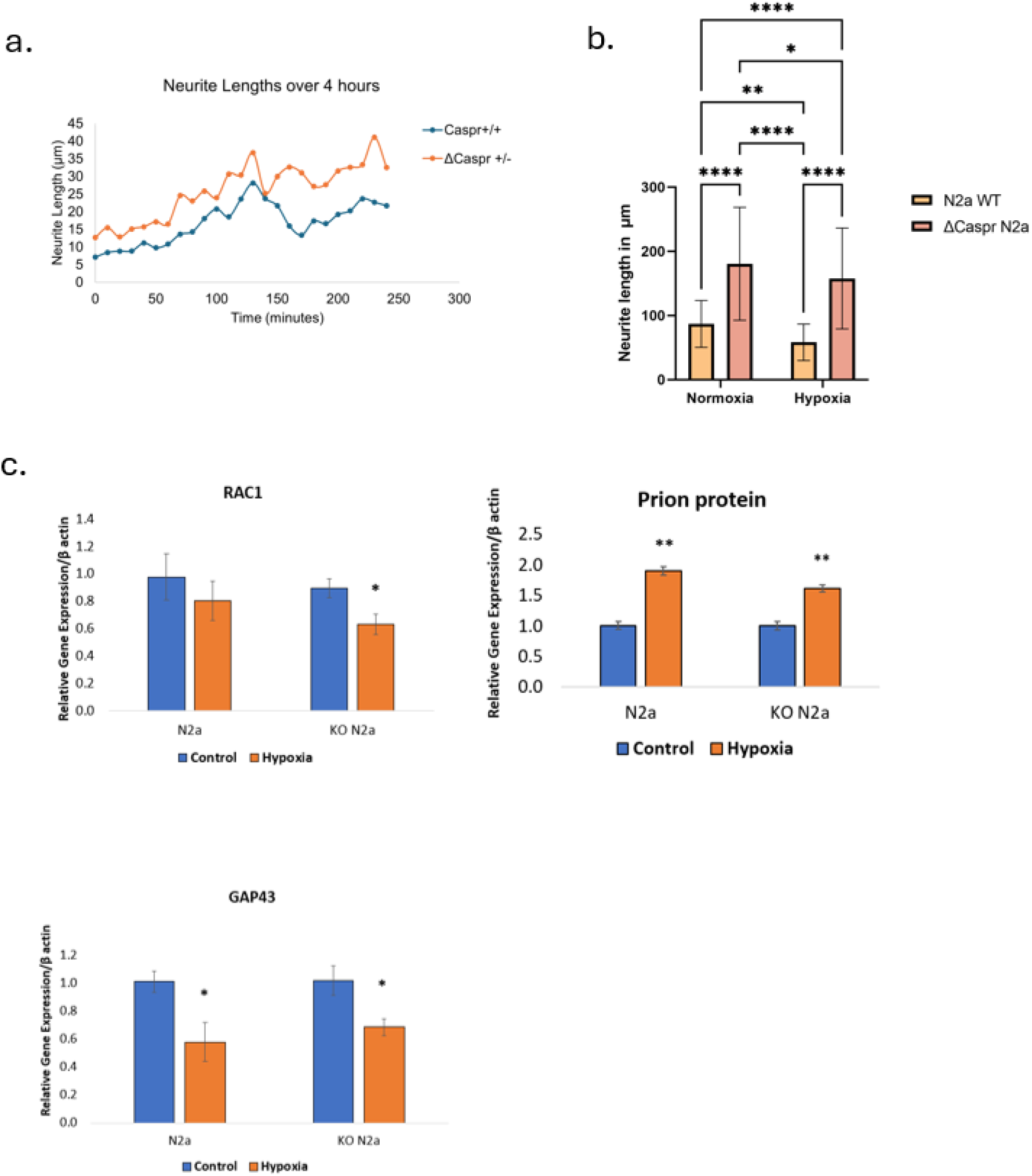
Impact of Caspr1 Deletion on Neurite Length and Neurite-Regulating Factors in Neuro2a Cells. This line graph illustrates the neurite length over a 4-hour period in Neuro2a cells, comparing wild-type (Caspr +/+) and knockout (Caspr +/-) phenotypes. The data underscore the critical role of Caspr1 in neurite extension and maintenance during neuritogenesis (N=3). A paired t-test was performed to compare the means of the two related groups. A one-tailed test was used to determine the statistical significance of the results. (b) The graph represents the neurite length (mean ± SEM) measured in wild-type (WT) and knockout (KO) cells under normoxia and hypoxia (N=3). Statistical analysis was performed using a two-way ANOVA to assess the main effects of cell type (WT vs. KO) and condition (normoxia vs. hypoxia), as well as their interaction. Tukey’s post hoc multiple comparisons test was applied to assess pairwise differences between each condition. Statistical significance is indicated by *p < 0.05, **p < 0.01 and ****p < 0.0001. (c) Bar graphs showing mean ± SEM of the relative gene expression levels of key neurite-regulating proteins—Prion protein, GAP43, and RAC1—in both wild-type (WT) and ΔCaspr1 Neuro2a cells under normoxic (control) and hypoxic conditions (N=3). The results provide insight into how the absence of Caspr1 affects the expression of these factors under different environmental stresses. An unpaired t-test (two-sample assuming equal variances) was performed to compare the groups. The degree of significance is represented as follows: **P<0.01 and *P<0.05.

### Hypoxia-Induced Alterations in Mitochondrial Morphology and Function

To investigate the effects of hypoxia on mitochondrial morphology, neurons were subjected to 24 hours of hypoxia, stained with TOM20 (a mitochondrial outer membrane marker), and analyzed via microscopy. Under normoxic conditions, mitochondria in neurons displayed typical tubular and branched structures. In contrast, hypoxia-treated neurons exhibited a shift towards more spherical mitochondrial morphology.

Western blot analysis of OPA1, a key regulator of mitochondrial fusion, revealed an increase in the short isoforms of OPA1 in hypoxia-treated neurons. This shift towards shorter OPA1 isoforms is indicative of enhanced mitochondrial fission and reduced fusion, leading to the observed fragmented mitochondrial structure.

To quantify mitochondrial sphericity, both wild-type and ΔCaspr1 Neuro2a cells were subjected to hypoxia, stained with MitoTracker Deep Red, and imaged. Mitochondrial sphericity was quantified using Imaris software, with results showing that hypoxia-treated cells had significantly increased mitochondrial sphericity compared to normoxia-treated cells. Notably, ΔCaspr1 cells exhibited higher mitochondrial sphericity under both normoxic and hypoxic conditions relative to wild-type cells, suggesting that Caspr1 plays a role in maintaining mitochondrial morphology.

Given these morphological changes, we further assessed the functional impact of hypoxia on mitochondria by measuring mitochondrial respiration rates using the Mito Stress assay on the Agilent Seahorse instrument. Cortical neurons were cultured in Seahorse 96-well plates, treated with 200 μM Cobalt Chloride (CoCl_2_) to mimic hypoxia for 24 hours, and then subjected to the assay. The results showed that hypoxia significantly impaired mitochondrial respiration, as evidenced by reductions in basal respiration, maximal respiration, ATP production rate, and spare respiratory capacity compared to neurons maintained under normoxic conditions.

These findings demonstrate that hypoxia not only induces significant changes in mitochondrial morphology, shifting towards a more spherical and fragmented structure, but also substantially impairs mitochondrial function, highlighting the detrimental effects of hypoxic stress on neuronal energy metabolism.

**Figure 6:**
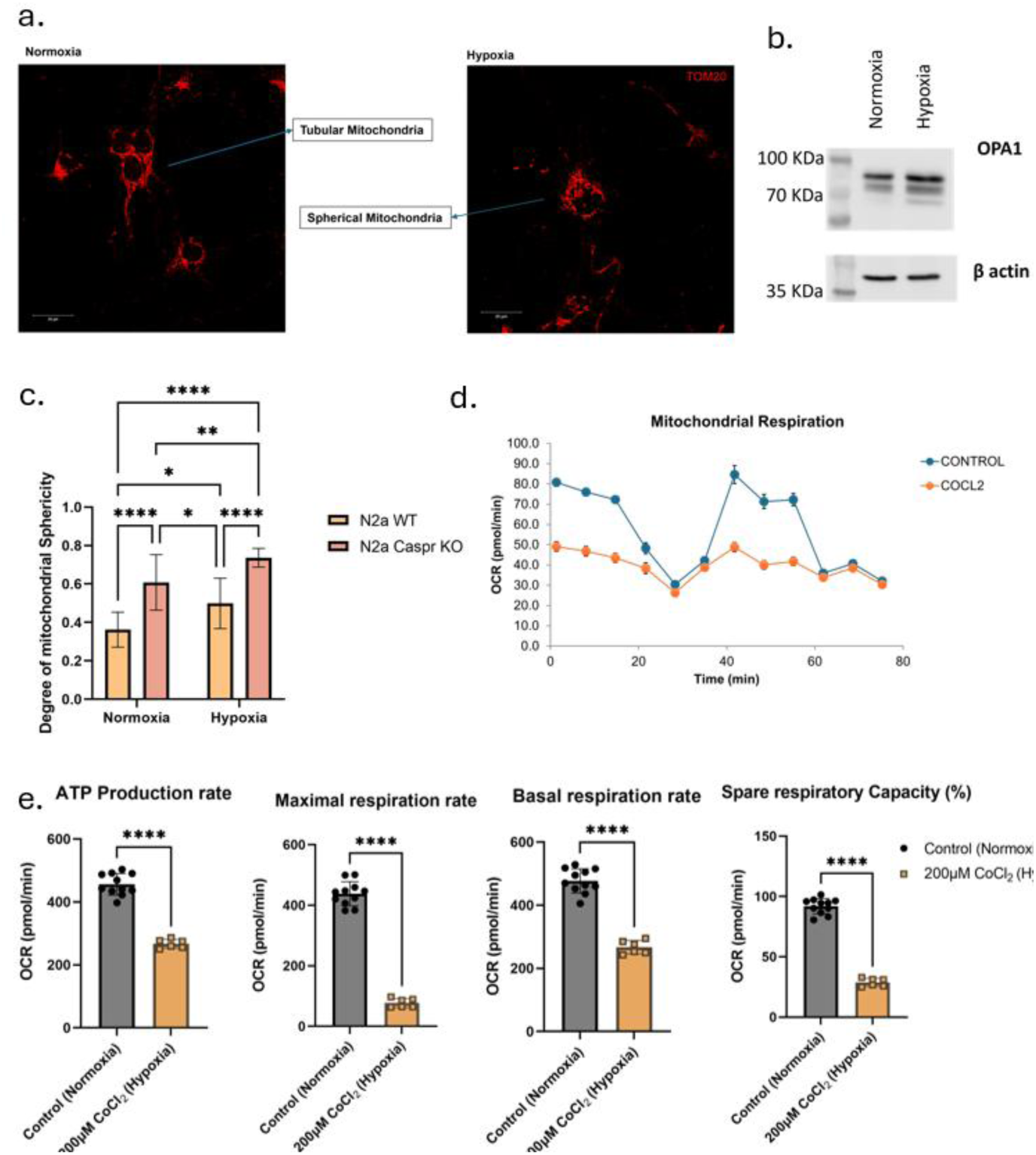
Effects of Hypoxia on Mitochondrial Morphology and Function in Neurons. (a) Immunofluorescence images of neurons stained with MitoTracker Deep Red, visualizing mitochondrial structure under normoxic and hypoxic conditions, and revealing hypoxia-induced morphological changes. (b) Western blot analysis of OPA1 protein levels in neurons under normoxic and hypoxic conditions, highlighting hypoxia’s impact on mitochondrial dynamics. (c) The graph represents the mitochondrial sphericity (mean ± SD) measured in N2a WT and ΔCaspr N2a cells under normoxia and hypoxia (N=3). Statistical analysis was performed using a two-way ANOVA to assess the main effects of cell type and condition (normoxia vs. hypoxia), as well as their interaction. Tukey’s post hoc multiple comparisons test was applied to assess pairwise differences between each condition. Statistical significance is indicated by *p < 0.05, **p < 0.01, ***p < 0.001 and ****p < 0.0001(d) A line graph depicting the mitochondrial respiration rate (measured as Oxygen Consumption Rate, OCR) in neuronal cells under normoxic and hypoxic (200 μM CoCl2) conditions, using the Agilent Seahorse instrument. Data represents as mean ± SEM (n=5) (e) Bar graphs presenting basal respiration, maximal respiration, ATP production rate, and spare respiratory capacity (%), illustrating the comprehensive impact of 24-hour hypoxia on mitochondrial respiratory function. Data is represented as mean ± SD from each calculation (n=5 technical replicates). An unpaired t-test was performed to compare the means of the two related groups. A two-tailed test was used to determine the statistical significance of the results. ****p<0.0001

## Discussion

Hypoxia is a well-documented factor in the pathophysiology of neurodegenerative diseases such as Alzheimer’s and Parkinson’s disease. For instance, studies have shown that hypoxic conditions can exacerbate the accumulation of amyloid-beta plaques in Alzheimer’s disease models, leading to increased neuronal death and cognitive decline (Tao et al., 2024). Similarly, in Parkinson’s disease models, hypoxia has been found to enhance the aggregation of alpha-synuclein, contributing to the degeneration of dopaminergic neurons (Gao et al., 2024).

In the context of brain injuries, hypoxia plays a critical role in the secondary injury cascade following traumatic brain injury (TBI). Research indicates that hypoxic conditions post-TBI can lead to increased neuronal apoptosis and neuroinflammation, further aggravating brain damage (Lindblad & Thelin, 2019). Additionally, stroke models frequently highlight the detrimental effects of hypoxia, where ischemic conditions result in rapid energy depletion and neuronal loss (Moskowitz et al., 2010).

Understanding the neuronal responses to hypoxic stress, as demonstrated in this study, is crucial for identifying potential therapeutic targets. For example, interventions aimed at modulating hypoxia-inducible factors (HIFs) or enhancing mitochondrial function may offer neuroprotective benefits in conditions characterized by hypoxic stress (Freeman & Barone, 2005; Mitroshina et al., 2021). Developing treatments that mitigate hypoxia-induced damage could significantly improve outcomes for patients with neurodegenerative diseases and brain injuries.

Our current study provides strong evidence for the role of Caspr in hypoxia-mediated neuritogenesis in embryonic cortical neurons, highlighting the interplay between hypoxic conditions, Caspr1 expression, transcriptional regulation of C/EBPα and mitochondrial dynamics. This is the first study to demonstrate this interconnection and the role of this paranodal protein in the hypoxia regulation of the neurons. Caspr is proven to be important for myelination and also has been shown as markers in specific diseases. Interestingly this heavily glycosylated, huge protein is expressed in different types of neurons and this may have neuron specific functions as well as the generic function of myelination. In another study from our lab submitted recently, Caspr is expressed in retinal neurons and regulates neuritogenesis under hyperglycemic condition, a direct implication in diabetes driven retinal complications (Data submitted for publication). Our results demonstrate that hypoxia significantly reduces neurite length while alternating between hypoxic and normoxic conditions effectively reverses this phenotype. This plasticity in neurons, which is normally not so plastic, suggests that neurons possess an adaptive response mechanism to environmental stress, which is critical for maintaining neuronal health and connectivity, failing which early onset of neurodegeneration can occur.

Caspr is increased in hypoxic conditions and as shown earlier from other studies including our own, Caspr inhibits neurite outgrowth. As an output, the neurite length measurement indicated a striking reduction in neurite length under hypoxic conditions. This outcome indicates that hypoxia adversely impacts neuronal growth and survival. The phenotypic reversibility of hypoxia-induced neurite length reduction upon switching to normoxia underscores the potential for neuronal recovery and adaptation. The significant increase in Caspr1 protein levels in hypoxic neurons also suggests that Caspr1 may play a protective role during hypoxic stress. The transcriptional regulation of Caspr1 by CEBPα adds an extra layer of complexity to this relationship, suggesting that CEBPα has a dual role, responding to hypoxic conditions and driving the expression of neurite regulating factors like Caspr1.

Growth cones are structures regulating the neurite elongation and driving synapse formation. Caspr 1 is expressed in growth cones. The observation of breaks in neurites and reduced growth cone size under prolonged hypoxic conditions emphasizes the detrimental effects of hypoxia on neuronal morphology. These morphological changes might also correlate with altered signaling dynamics in growth cones, warranting further investigation into the signaling pathways involved. The temporal analysis of Caspr1 expression revealed a nuanced response to hypoxia, with initial downregulation at early time points followed by significant upregulation at 12 and 24 hours. This dynamic regulation suggests that Caspr1 may initially serve as a stress response factor that becomes increasingly important as hypoxic conditions persist. Such a time-dependent response could have implications for therapeutic timing in interventions aimed at promoting neuronal survival. We also examined other neurite-regulating proteins, such as Prion protein and RhoA, whose expression also increases in hypoxia conditions. The Prion protein is an established neurite-promoting molecule similar to Rho A. Our data shows that, even when the Prion protein is increasing, the neurite outgrowth is reduced, suggesting the dominance rendered by reduced Caspr expression.

Interestingly, ΔCaspr1 Neuro2a cells exhibited significantly longer neurites under hypoxic conditions compared to their wild-type counterparts, suggesting that the loss of Caspr1 may confer a degree of resistance to hypoxia-induced morphological changes. This finding highlights a potential compensatory mechanism in the absence of Caspr1 and suggests that targeting Caspr1 pathways could enhance neuronal resilience during hypoxic events.

Our investigation into mitochondrial morphology revealed a shift from tubular and branched structures to more spherical forms under hypoxia, along with an increase in short isoforms of OPA1, which helps in maintaining balance between fission and fusion of mitochondria. This elucidates that there is an enhanced mitochondrial fission, associated with impaired mitochondrial function. This is further being investigated in the lab. Subsequently, we assessed mitochondrial respiration rates which revealed significant declines under hypoxia, clearly indicating hypoxia not only alters mitochondrial morphology but also impairs their functional capacity, a parameter that needs further elucidation. The observed relationship between the expression of Caspr1 and mitochondrial dynamics further emphasizes the need to understand how Caspr1 influences mitochondrial morphology and function under stress conditions. ΔCaspr1 cells displayed higher mitochondrial sphericity, suggesting that Caspr1 may also play a role in maintaining mitochondrial integrity.

In summary, this study highlights the critical role of Caspr in the response of cortical neurons to hypoxia, detailing how its expression is regulated by hypoxic stress and its subsequent effects on neurite morphology and mitochondrial dynamics. The findings suggest that Caspr1 is an important player in neuronal plasticity and resilience, with potential implications for therapeutic strategies aimed at mitigating the effects of hypoxia in neurodegenerative conditions. The changes in expression patterns of these proteins under hypoxic conditions suggest a comprehensive reprogramming of neurite growth mechanisms, highlighting the complexity of neuronal responses to oxygen deprivation. Deeper studies in this direction and the connection between hypoxia in humans may be understood by studying populations living in high altitudes and soldiers posted in high altitudes, an initiative that we are pursuing in the lab currently. Future studies from the lab should aim to unravel the specific signaling pathways involved in Caspr1 regulation and explore potential interventions that target these pathways to enhance neuronal survival in hypoxic environments. Since Caspr 1 and CEBPα are elevated in hypoxia-treated neurons, it may be considered as a potential therapeutic target for enhancing neuronal resilience.

## References

Al Tameemi, W., Dale, T. P., Al-Jumaily, R. M. K., & Forsyth, N. R. (2019). Hypoxia-Modified Cancer Cell Metabolism. Frontiers in Cell and Developmental Biology, 7. 10.3389/FCELL.2019.00004

Anderson, G. R., Galfin, T., Xu, W., Aoto, J., Malenka, R. C., & Südhof, T. C. (n.d.). Candidate autism gene screen identifies critical role for cell-adhesion molecule CASPR2 in dendritic arborization and spine development. 10.1073/pnas.1216398109

Bennison, S. A., Blazejewski, S. M., Smith, T. H., & Toyo-Oka, K. (2020). Protein kinases: master regulators of neuritogenesis and therapeutic targets for axon regeneration. Cellular and Molecular Life Sciences: CMLS, 77(8), 1511–1530. 10.1007/s00018-019-03336-6

Coman, I., Aigrot, M. S., Seilhean, D., Reynolds, R., Girault, J. A., Zalc, B., & Lubetzki, C. (2006). Nodal, paranodal and juxtaparanodal axonal proteins during demyelination and remyelination in multiple sclerosis. Brain, 129(12), 3186–3195. 10.1093/BRAIN/AWL144

Devanathan, V., Jakovcevski, I., Santuccione, A., Li, S., Lee, H. J., Peles, E., Leshchyns’ka, I., Sytnyk, V., & Schachner, M. (2010). Cellular Form of Prion Protein Inhibits Reelin-Mediated Shedding of Caspr from the Neuronal Cell Surface to Potentiate Caspr-Mediated Inhibition of Neurite Outgrowth. Journal of Neuroscience, 30(27), 9292– 9305. 10.1523/JNEUROSCI.5657-09.2010

Einheber, S., Zanazzi, G., Ching, W., Scherer, S., Milner, T. A., Peles, E., & Salzer, J. L. (1997). The axonal membrane protein Caspr, a homologue of neurexin IV, is a component of the septate-like paranodal junctions that assemble during myelination. The Journal of Cell Biology, 139(6), 1495–1506. 10.1083/JCB.139.6.1495

Falivelli, G., De Jaco, A., Lietta Favaloro, F., Kim, H., Wilson, J., Dubi, N., Ellisman, M. H., Abrahams, B. S., Taylor, P., & Comoletti, D. (n.d.). Inherited genetic variants in autism-related CNTNAP2 show perturbed trafficking and ATF6 activation. 10.1093/hmg/dds320

Fernandes, D., Santos, S. D., Coutinho, E., Whitt, J. L., Beltrão, N., Rondão, T., Leite, M. I., Buckley, C., Lee, H.-K., & Carvalho, A. L. (n.d.). Disrupted AMPA Receptor Function upon Genetic-or Antibody-Mediated Loss of Autism-Associated CASPR2. 10.1093/cercor/bhz

Freeman, R. S., & Barone, M. C. (2005). Targeting hypoxia-inducible factor (HIF) as a therapeutic strategy for CNS disorders. Current Drug Targets. CNS and Neurological Disorders, 4(1), 85–92. 10.2174/1568007053005154

Gao, Y., Zhang, J., Tang, T., & Liu, Z. (2024). Hypoxia Pathways in Parkinson’s Disease: From Pathogenesis to Therapeutic Targets. International Journal of Molecular Sciences, 25(19). 10.3390/IJMS251910484

H, F., AC, Z., J, X., A, M., D, Z., EY, S., & GG, H. (2011). Severe Hypoxia: Consequences to Neural Stem Cells and Neurons. Journal of Neurology Research, 1(5). 10.4021/JNR70W

Hofmeijer, J., Mulder, A. T. B., Farinha, A. C., Van Putten, M. J. A. M., & Le Feber, J. (2014). Mild hypoxia affects synaptic connectivity in cultured neuronal networks. Brain Research, 1557, 180–189. 10.1016/J.BRAINRES.2014.02.027

Kaelin, W. G., & Ratcliffe, P. J. (2008). Oxygen sensing by metazoans: the central role of the HIF hydroxylase pathway. Molecules and Cells, 30(4), 393–402. 10.1016/J.MOLCEL.2008.04.009

Lindblad, C., & Thelin, E. P. (2019). Secondary Insults in Experimental Traumatic Brain Injury: The Addition of Hypoxia. Neuromethods, 149, 223–242. 10.1007/978-1-4939-9711-4_13

Mitroshina, E. V., Savyuk, M. O., Ponimaskin, E., & Vedunova, M. V. (2021). Hypoxia-Inducible Factor (HIF) in Ischemic Stroke and Neurodegenerative Disease. Frontiers in Cell and Developmental Biology, 9. 10.3389/FCELL.2021.703084/PDF

Moskowitz, M. A., Lo, E. H., & Iadecola, C. (2010). The science of stroke: mechanisms in search of treatments. Neuron, 67(2), 181–198. 10.1016/J.NEURON.2010.07.002

Nakayama, K., & Kataoka, N. (2019). Regulation of Gene Expression under Hypoxic Conditions. International Journal of Molecular Sciences, 20(13), 3278. 10.3390/IJMS20133278

O’Driscoll, C. M., & Gorman, A. M. (2005). Hypoxia induces neurite outgrowth in PC12 cells that is mediated through adenosine A2A receptors. Neuroscience, 131(2), 321–329. 10.1016/J.NEUROSCIENCE.2004.11.015

Pocock, R., & Hobert, O. (2008). Oxygen levels affect axon guidance and neuronal migration in Caenorhabditis elegans. Nature Neuroscience 2008 11:8, 11(8), 894–900. 10.1038/nn.2152

Saha, D., Vishwakarma, S., Gupta, R. K., Pant, A., Dhyani, V., Sharma, S., Majumdar, S., Kaur, I., & Giri, L. (2023). Non-prophylactic resveratrol-mediated protection of neurite integrity under chronic hypoxia is associated with reduction of Cav1.2 channel expression and calcium overloading. Neurochemistry International, 164. 10.1016/J.NEUINT.2022.105466

Schröder, R., & Luhmann, H. (1997). Morphology, Electrophysiology and Pathophysiology of Supragranular Neurons in Rat Primary Somatosensory Cortex. European Journal of Neuroscience, 9(1), 163–176. 10.1111/J.1460-9568.1997.TB01364.X

Su, K. (1975). Brain hypoxia studied in mouse central nervous system cultures. I. Sequential cellular changes. Laboratory Investigation.

Tao, B., Gong, W., Xu, C., Ma, Z., Mei, J., & Chen, M. (2024). The relationship between hypoxia and Alzheimer’s disease: an updated review. Frontiers in Aging Neuroscience, 16, 1402774. 10.3389/FNAGI.2024.1402774/BIBTEX

Wallace, M. G., Hartle, K. D., Snow, W. M., Ward, N. L., & Ivanco, T. L. (2007). Effect of hypoxia on the morphology of mouse striatal neurons. Neuroscience, 147(1), 90–96. 10.1016/j.neuroscience.2007.02.067

Wolswijk, G., & Balesar, R. (2003). Changes in the’expression and localization of the paranodal protein Caspr on axons in chronic multiple sclerosis. Brain, 126(7), 1638– 1649. 10.1093/BRAIN/AWG151

Xiong, T., Yang, X., Qu, Y., Chen, H., Yue, Y., Wang, H., Zhao, F., Li, S., Zou, R., Zhang, L., & Mu, D. (2019). Erythropoietin induces synaptogenesis and neurite repair after hypoxia ischemia-mediated brain injury in neonatal rats. NeuroReport, 30(11), 783–789. 10.1097/WNR.0000000000001285

